# Virulence of *Naegleria fowleri* isolates varies significantly in the mouse model of primary amoebic meningoencephalitis

**DOI:** 10.1101/2025.05.20.655168

**Authors:** A. Cassiopeia Russell, Erica Schiff, Joseph Dainis, Håkon Jones, Christopher A. Rice, Dennis E. Kyle

## Abstract

*Naegleria fowleri* is a small free-living amoeba that causes an acute, fatal disease called primary amoebic meningoencephalitis (PAM). One persisting question is why few people succumb to disease when so many are potentially exposed. We tested the hypothesis that *N. fowleri* isolates vary in virulence and in the minimum infectious dose required to induce disease by using a mouse model of PAM and intranasally inoculating dilutions of five clinical isolates of *N. fowleri.* Results showed significant differences in onset of severe disease and mortality rates between isolates. Remarkably, for isolate V596, 100 amoebae produced 100% mortality within 5 days. In contrast, higher numbers of amoebae were required for other isolates and mice survived for >2 weeks. Concurrently, we developed an *in vitro* virulence assay by comparing feeding rates between amoebae isolates seeded onto Vero cells. We observed a positive correlation between cytopathic effects *in vitro* and virulence *in vivo*.

**Article Summary Line:** Isolates of *Naegleria fowleri* vary in virulence, which results in significant differences in minimal infectious concentrations of amoebae that cause primary amoebic meningoencephalitis.

---

Small, thermotolerant free-living amoebae are ubiquitous in soil and freshwater in the environment, although several pathogenic free-living amoebae are known to cause serious diseases in humans.(1) *Naegleria fowleri*, colloquially known as the brain-eating amoeba, is the causative agent of primary amoebic meningoencephalitis (PAM). Humans become infected while participating in freshwater activities or performing nasal ablutions when water containing the amoebae enters the nasal passages; the amoebae then traverse the olfactory epithelium and migrate to the frontal lobes of the brain where they cause significant pathology. Meningitis symptoms usually begin within a week after warm water exposure, and severe complications followed by death occur in the next 1-18 days. Although PAM reportedly is a rare disease, it unfortunately has a >98% case fatality rate with no survivors in the US since 2016.(2) There are several lines of evidence that suggest PAM is under-reported in warm climates around the world. Firstly, in Pakistan, where ritualistic nasal ablution is a common practice, large outbreaks have been documented recently once surveillance was instituted.(3, 4) In addition, the early signs and symptoms of PAM are similar to meningitides caused by viral or bacterial pathogens, therefore most PAM cases are misdiagnosed or detected late in the infection, or post- mortem.(1) Lastly, recent studies reported high rates of seropositivity to *N. fowleri* in people from several regions of Mexico.(5) These data suggest high rates of environmental exposure to *N. fowleri*, which contrasts sharply with the low rates of infection and disease in humans.

Quite often the source of infection with *N. fowleri* is linked to a site where hundreds of people presumably had similar exposure to amoebae in the same time frame as the individual that contracted the deadly infection. These include lakes,(6, 7) hot spring-fed pools,(8) whitewater recreation areas,(9) inadequately treated potable water (e.g., non-sterile tap water used in neti-pots),(10) and artificial water sports venues,(11) to name a few. Since the numbers of documented PAM cases are low and exposure to amoebae in warm water is high, a major question is why some people get infected with *N. fowleri* and develop PAM, whereas others do not. There are many possible hypotheses for low rates of infection and disease; these include host and parasite factors which are difficult to study with the currently available tools.

In this study we hypothesized that *N. fowleri* isolates vary in virulence and that these differences affect the minimal infectious concentrations of amoebae required to initiate a fulminant infection. To test our hypothesis, we evaluated virulence of five clinical isolates of *N. fowleri* in the mouse model of PAM. In parallel, we also developed an *in vitro* cytopathogenicity model that demonstrated positive correlation between *in vivo* and *in vitro* virulence phenotypes.

## Methods

### N. fowleri isolates

All isolates of *N. fowleri* used in this study were collected from clinical cases of PAM (Table 1). ATCC 30215 (Nf69) was purchased from the American Type Culture Collection (ATCC), and the remaining isolates were generously provided by Dr. Ibne Ali, US Centers for Disease Control. The dates of isolation for the cases ranged from 1966 – 2011 and most of the patients were male (10 of 13). The isolates included examples of three genotypes, although genotypes I and III were the best represented groups in this collection. All isolates were routinely maintained in axenic culture in Nelson’s complete media (NCM) as previously described (12).

**Table 1.** List of *Naegleria fowleri* strains used in this study.

| Isolate | Sex/Location/Year | Genotype |
| --- | --- | --- |
| HB1 | M/Florida/1966 | III |
| Nf69* | M/Australia/1969 | IV |
| TY | M/Virginia/1969 | III |
| HB4 | F/Virginia/1977 | III |
| V067* | M/Arizona/1987 | III |
| V206 | M/Mexico/1990 | I |
| Villa Jose* | F/California/1996 | I |
| V413 | M/Texas/1998 | I |
| Davis | M/Florida/1998 | I |
| V455 | M/Nevada/2000 | III |
| V596* | M/Nevada/2007 | III |
| V631* | M/Louisiana | I |

\* Used for *in vivo* studies

### *In vivo* virulence

In preparation for an *in vivo* experiment, first the amoebae from axenic cultures were centrifuged and washed 2 times in 1X phosphate buffered saline (PBS) before adding to flasks containing Vero cell (green monkey kidney cells; E6; ATCC CRL-1586) monolayers (Figure 1). Vero cells were cultured and sub-cultured as previously described (13). The amoebae-host cell cocultures were monitored daily and subculture to a new passage was made once most (≥85%) of the Vero cells had been destroyed by the amoebae. We passaged the amoebae ≥ 6 times on Vero cells to “prime” them for infection and then collected and concentrated amoebae by centrifugation before preparing stock concentrations containing 1000, 10,000, or 50,000 trophozoites in 100 µl of PBS. Then 10 µl of PBS containing amoebae was then inoculated into a single nare of an anesthetized female 3–4-week-old ICR (CD-1) mouse, ranging from 17-26.5g in weight. Either 5 or 8 mice were infected per isolate-concentration study, whereas n=5 mice were used as uninfected control animals.

**Figure 1.**
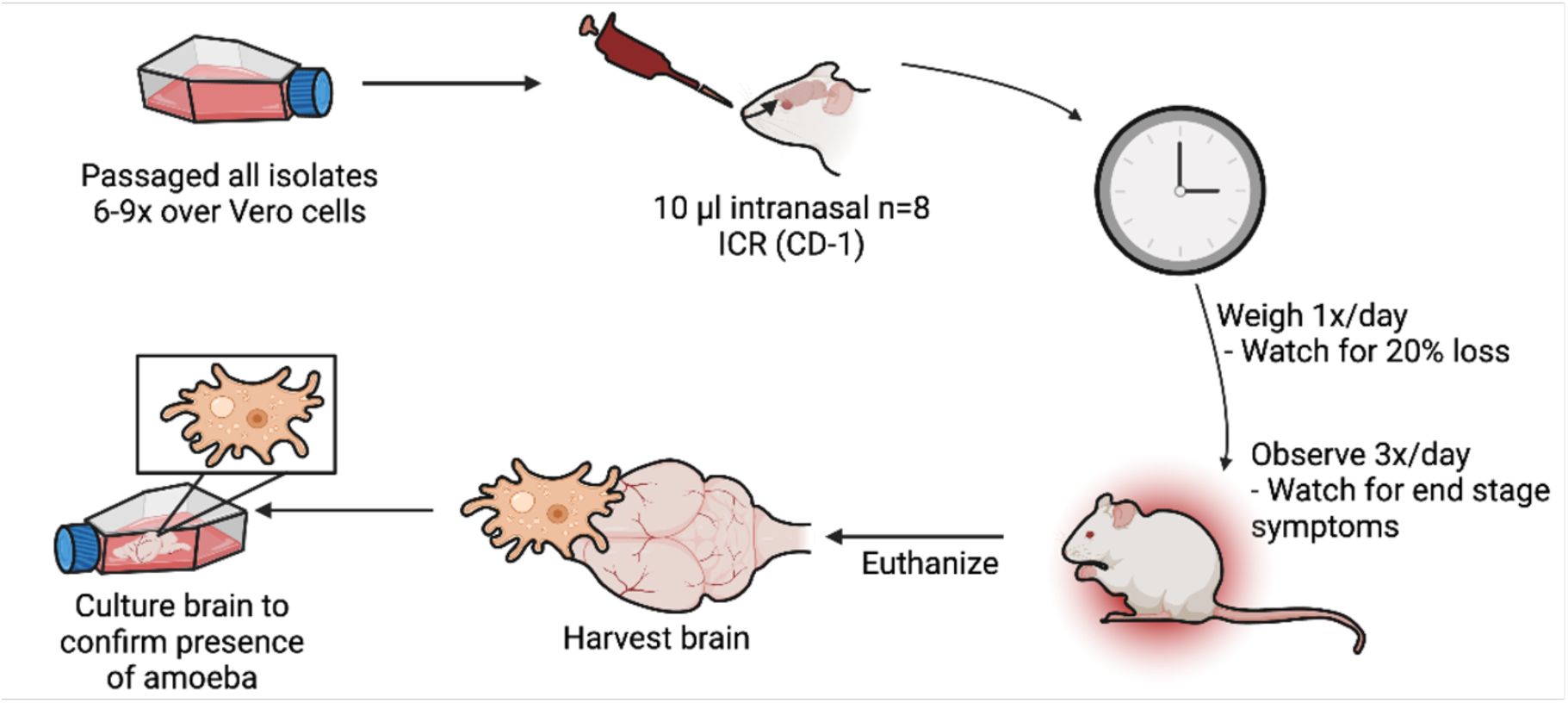
Schematic overview of methods used for *in vivo* virulence studies. *N. fowleri* strains were first passaged over Vero cells at least 6 times. Amoebae were washed and diluted to the desired amoeba number in 10 μL and then dosed intranasally into mice (n=8 per group). Animals were observed multiple times per day and euthanized when symptoms of disease were observed. Brains of mice were harvested and inoculated into culture media to confirm the presence of amoebae. (Created with BioRender.com.)

Daily weight monitoring was performed as well as thrice daily inspections for symptoms of PAM, including piloerection, orbital tightening, arched spine, ataxia, increased respiratory effort and seizures. Animals displaying ≥20% weight loss or end-stage symptoms were euthanized per approved animal use protocols to mitigate pain or suffering. Brains were dissected from each animal; each brain was then placed into a flask containing 10 ml of NCM containing 10 U/ml of penicillin and streptomycin, and cultured for up to 7 days to microscopically confirm presence of amoebae. All surviving mice were euthanized on day 35 post infection to conclude the study.

An additional follow-up animal study was performed as described above to identify a minimum infectious dose for isolates that caused acute mortality at the lowest doses of 100 and 1000 amoebae. We utilized n=5 mice per concentration of isolate, with n=3 uninfected control mice. Isolate V596 was inoculated at concentrations of amoebae of 1000, 100, 75, 50, 25, or 10 per mouse. Isolate Villa Jose was inoculated at concentrations of amoebae of 5000, 2500, 1000, 500, 250, or 100 per mouse. Data from these studies were used to determine the 50% lethal concentrations (LC_50_s) of *N. fowleri* amoebae required to induce a fulminant infection in mice.

Survival data and LC_50_s were calculated by using GraphPad Prism software (version 10, GraphPad, La Jolla, CA, USA). All animal studies were reviewed and approved by the University of Georgia Institutional Animal Care and Use Committee (Protocol A202 03-026). All animal studies were conducted according to the Guide for the Care and Use of Laboratory Animals.

### *In vitro* virulence

For the *in vitro* virulence assay, we used the culture methods described above and first passaged each isolate 5 times over Vero cell monolayers in culture flasks. We then concentrated the cells by centrifugation and split the amoebae into three flasks for biological replicates. To prepare for the assay, 24 hours before the 5^th^ passage of amoebae finished feeding upon monolayers, we added 25,000 Vero cells/well in a µClear black Cellstar 96-well microplate (Greiner Bio-One, Kremsmünster, Austria) and incubated for two days at 37°C until the monolayer reached 80-90% confluency was formed. For initial optimization, test plates contained a dilution series of 100 to 50,000 amoebae and time checks were performed at 3 h, 6 h, 12 h, and 24 h to determine ideal amoebae seeding density and time for the cytopathogenicity assay. We determined that a maximal and minimal concentration of 10,000 and 625 amoebae, respectively, and an incubation time of 24h allowed for visible clearance of monolayers that could be measured with high content imaging. For the assay, passage 5 amoebae were harvested and serial dilutions of amoebae, from 10,000 to 625, were prepared and each concentration was added in quadruplicate wells containing confluent Vero cells.

After 24 h co-incubation, media was carefully aspirated to avoid disturbing adherent cells, and 100 µl of 4% paraformaldehyde (PFA) in phosphate-buffered saline (PBS) was added with an incubation period of 15 min in the dark at room temperature. Fixative was removed and a wash with 100 µl PBS was performed prior to adding 100 µl of 10 µg/ml Hoechst 33342. Plates were incubated at room temperature in the dark for 45 min prior to stain removal and a final PBS wash. PBS (100 µl) was added to each well and high content imaging was conducted on an ImageXpress Micro Confocal HCI system (Molecular Devices, San Jose, CA, USA) to identify nuclei stained with Hoechst 33342, quantify host cell count, and exclude smaller amoebae nuclei from cell totals. Final imaging parameters were set to count Vero cell nuclei, and the assay endpoint was the number of Vero cell nuclei remaining after 24 h. Vero-cell-only control wells were compared to amoeba serial dilution wells and percentage feeding rate comparisons were prepared using GraphPad Prism software (version 10, GraphPad, La Jolla, CA, USA).

## Results

Our first aim was to directly compare levels of virulence of different clinical isolates of *N. fowleri* in the ICR (CD-1) mouse model of PAM. For these studies we used five clinical isolates (Table 1). Given the lack of information regarding prior passage number and culture conditions of the isolates, we devised a scheme to allow direct comparison of these isolates. Previous studies suggest that passaging amoebae over mammalian cells can enhance pathogenicity in mouse infections.(14) Therefore, we standardized the preparation of amoebae for animal studies by passaging them 6-9 times over Vero cells to induce active feeding before preparing dilutions to be used to initiate PAM infection in mice (Figure 1).

The utility of this approach was demonstrated with a comparison of virulence in mice that were inoculated with amoebae from standard axenic culture (Nelson’s media) versus amoebae passaged over Vero cells. As shown in Figure 2, both the V596 and Nf69 strains of *N. fowleri* that were fed on Vero cells were significantly more virulent in mice than amoebae grown in axenic media. In addition, the enhanced virulence phenotype was observed with the most virulent strain (V596) as well as with the less virulent Nf69 strain.

**Figure 2.**
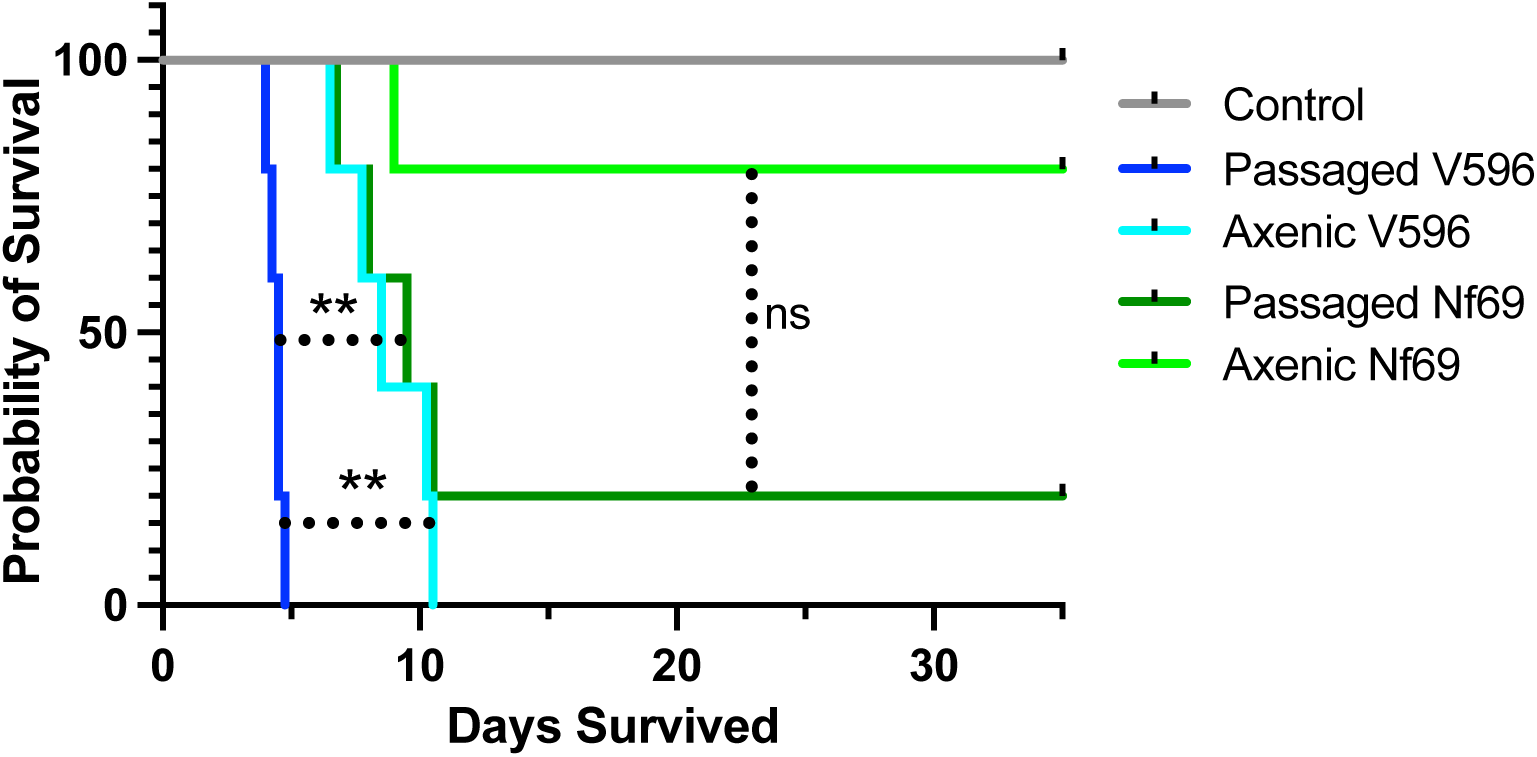
Survival curves for mice infected with lowly (Nf69) and highly (V596) virulent *N. fowleri* isolates show significant differences in survival. Differences in survival are also noted for passaged vs axenically cultured isolates (\*\**p* < 0.01; ns = not significant).

After validating the method, next we assessed virulence of five isolates of *N. fowleri* in the mouse model of PAM. Interestingly, these strains demonstrated significantly different levels of virulence *in vivo*, with mean survival times ranging from 4 days to ≥35 days (Figure 3). Nf69, a strain we have used as a lab standard for drug discovery studies,(15, 16) was moderately virulent. All mice inoculated with 5000 Nf69 amoebae succumbed to infection, yet only 75% and 25% died after infecting mice with 1000 or 100 amoebae, respectively. In contrast, V596 was the most virulent isolate tested. We observed 100% mortality with a mean survival time of 4-5 days, even with as few as 100 amoebae inoculated into a single nare of a mouse. V631, Villa Jose, and V067 produced mean survival times ranging from 6 to 14 days with inocula of 1000 amoebae per mouse. Interestingly, 25% of the mice succumbed to disease that were infected with 100 Villa Jose or V631; all mice inoculated with 100 V067 amoebae survived for 35 days. Differences in gross pathology of brains were observed in mice infected with V596 versus less virulent N. fowleri strains (Appendix Figure 1). These data demonstrate significant differences in virulence of *N. fowleri* strains.

**Figure 3.**
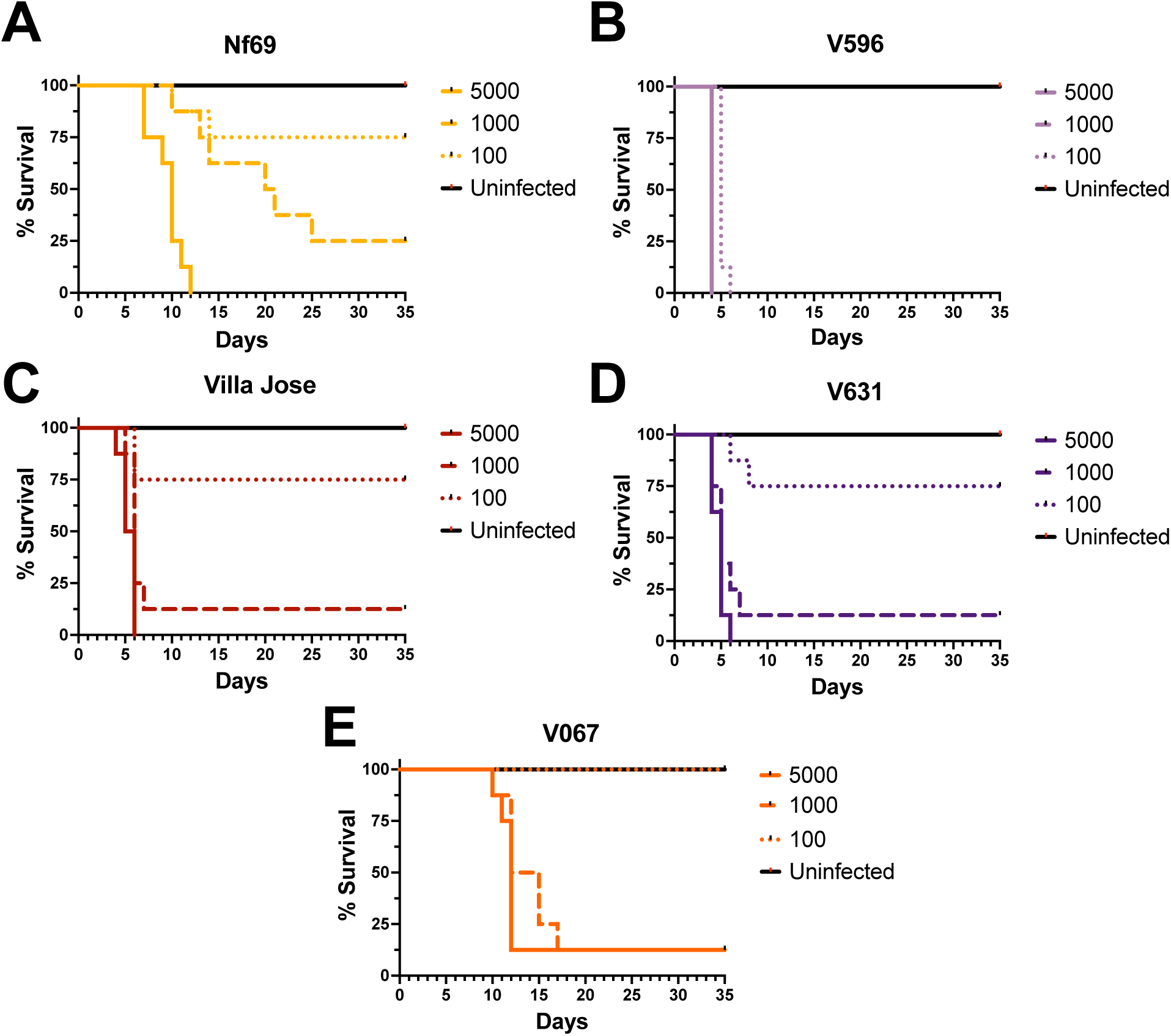
*N. fowleri* isolates exhibit varying levels of virulence in the mouse model of Primary Amoebic Meningoencephalitis (PAM). Amoeba numbers inoculated intranasally for each strain were 100, 1000, and 5000. The survival times to euthanasia due to severe symptoms of disease are shown for Nf69 (A), V596 (B), Villa Jose (C), V631 (D), and V067 (E), respectively.

In follow up mouse studies we conducted a comparison study to assess the 50% lethal concentrations of V596 and Villa Jose amoebae to induce a fatal PAM disease (Figure 4). For these studies we inoculated mice with to 10, 25, 50, 75, 100, and 1000 amoebae and assessed mouse survival times. These studies confirmed that V596 is the most virulent isolate tested, with deaths observed in <6 days with as few as 10 – 25 amoebae. The survival curves for Villa Jose also were reproducible with inocula of <u>≤</u>500 amoebae producing fulminant infections.

**Figure 4.**
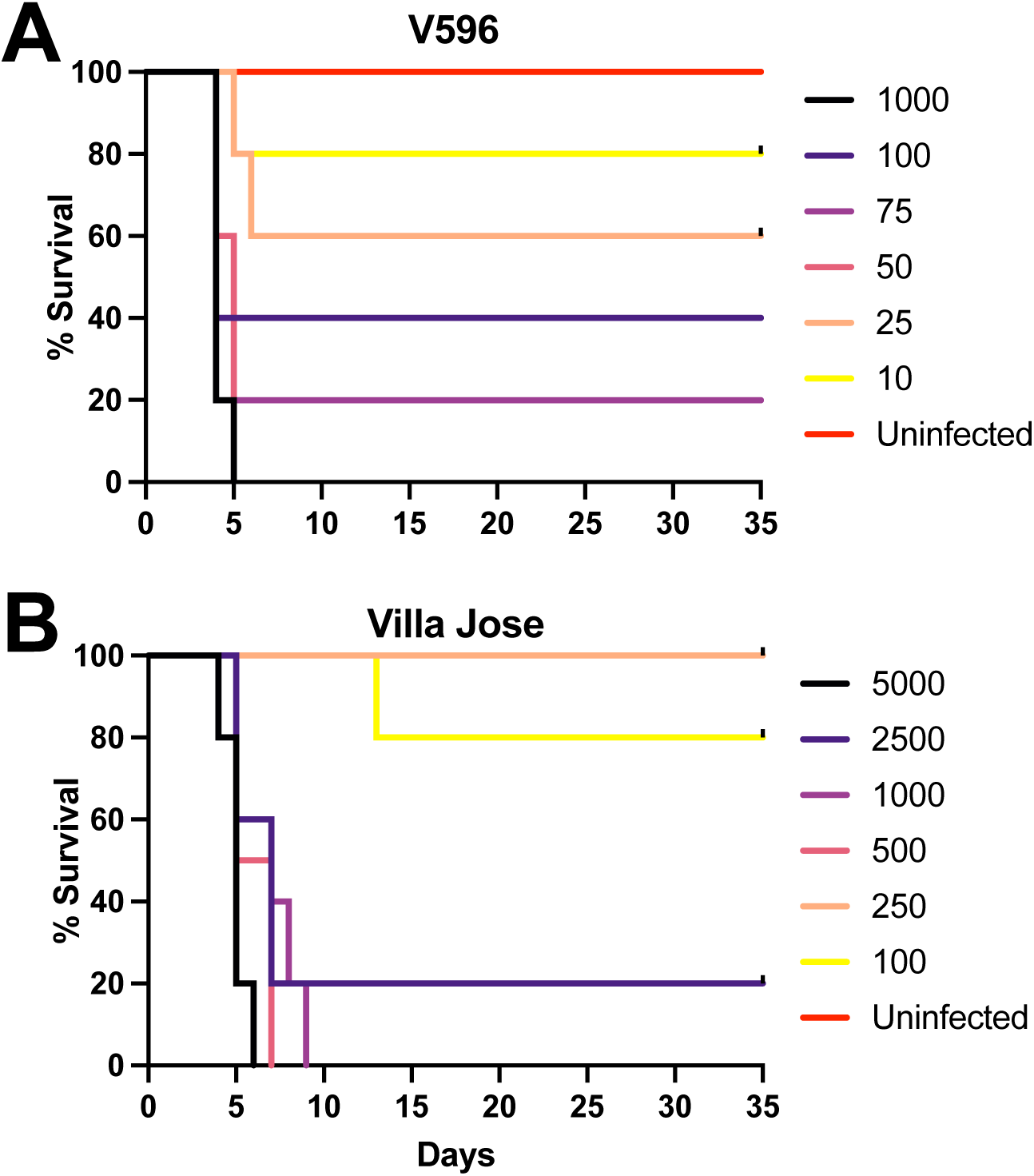
A follow-up *in* vivo study was conducted to identify the minimal infectious concentrations of V596 (A) and Villa Jose (B) strains of *N. fowleri* amoebae that caused 100% mortality in the mouse model of PAM. V596 was found to be the most virulent strain, requiring only 10 amoebae to cause death and an estimated 50% lethal concentration of ∼40 amoebae. The minimal 100% lethal concentration of Villa Jose was 1000 amoebae, with an estimated LC_50_ of 325 amoebae.

Based upon these studies, the estimated 50% lethal concentrations (LC50s) are ∼40 and ∼325 amoebae for V596 and Villa Jose, respectively. In addition to providing quantitative estimates of virulence differences between isolates, the reproducibility of survival data demonstrates the utility of the standardized method of feeding amoebae over Vero cells prior to conducting *in vivo* studies.

### *In vitro* virulence assay

For additional studies on mechanisms of virulence in *N. fowleri*, it would be advantageous to have a facile, quantitative *in vitro* assay for assessing virulence phenotype:genotype associations in a high-throughput assay. During the conduct of the Vero cell passages for the *in vivo* studies, we observed that isolates cleared the Vero cell monolayers at different rates. Some isolates would destroy the complete Vero cell monolayer within 24 h, whereas others would take several days to clear the mammalian cells. Thus, we hypothesized that the rate at which *N. fowleri* isolates feed on Vero cells could serve as a surrogate endpoint for virulence, with the additional prospect of serving as a surrogate of *in vivo* virulence.

For the *in vitro* virulence assay, we prepared the amoebae by passaging them over Vero cells ≥5 times before diluting the amoebae for initiation of the virulence assays. In preliminary studies we tested different time points (24-120 h) and the endpoint for host cell killing was determined by high-content imaging of Hoechst 33342 stained nuclei. Upon assessing the cytopathic effects as shown in Figure 5, we observed significant differences in clearance of Vero cells from co-cultures with 12 isolates of *N. fowleri* (Figure 6). The most virulent isolates were V596, V631, Villa Jose, and V206, with > 99% of the Vero cells cleared by 10,000 amoebae within 24 h. Conversely, the least virulent strains cleared <5% of Vero cells from the cocultures in 24 h. Some *N. fowleri* strains were moderately virulent with feeding rates varying between 5- 50% within 24 h. Of the 12 isolates assessed in the in vitro assay, five were genotype I, and six were genotype III. We observed no correlation between virulence *in vitro* and these genotypes (Appendix Figure 2). Isolates from both genotypes I and III produced a wide spectrum of virulence *in vitro*, suggesting strain dependent factors are responsible for virulence. Results from the *in vitro* and *in vivo* assays demonstrated a positive correlation of virulence in the two models tested with five *N. fowleri* strains (Figure 7).

**Figure 5.**
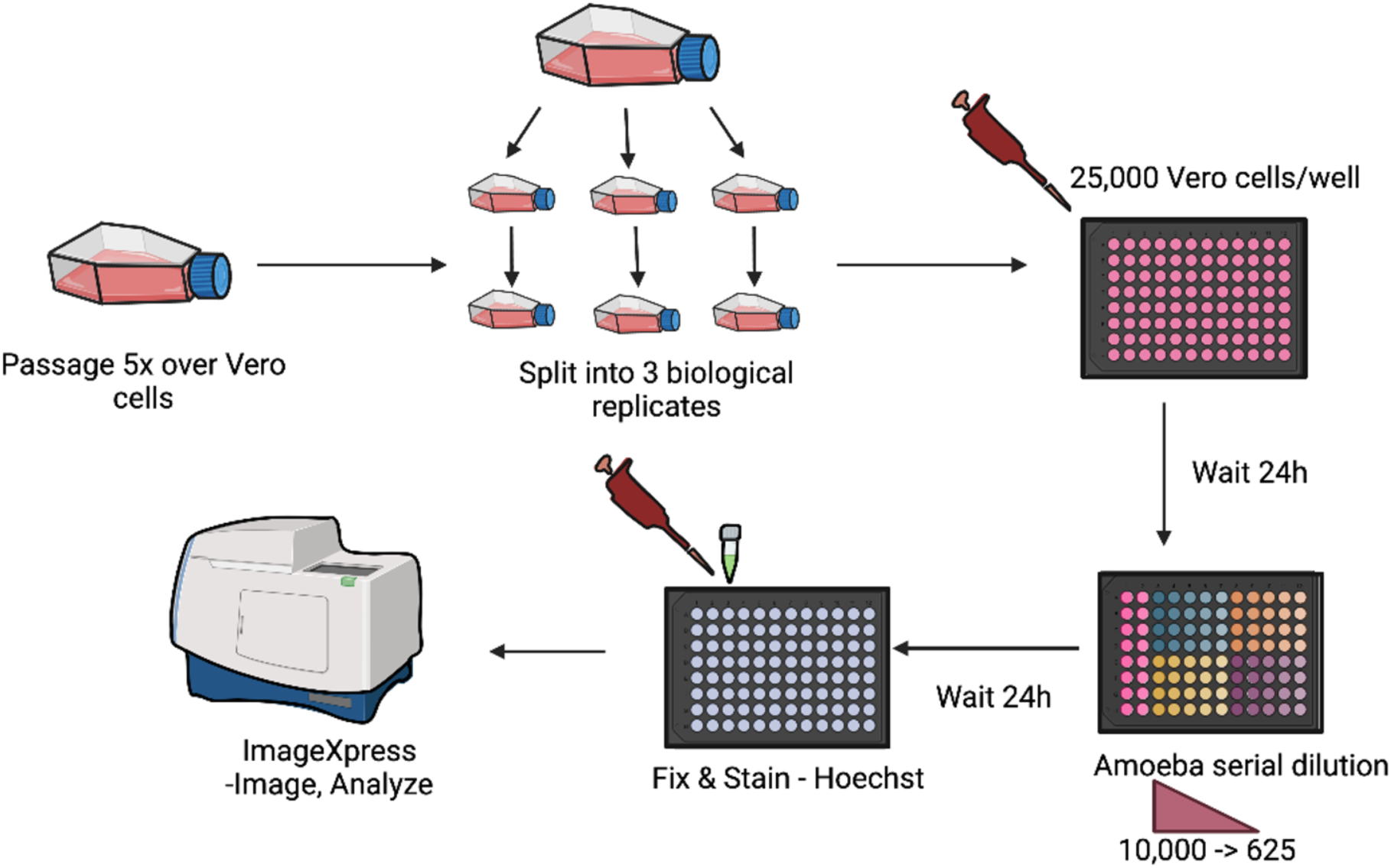
Schematic of methods used for the in vitro virulence assay. *N. fowleri* trophozoites were passaged 5 times over Vero cells, washed, and then split into three biological replicates. Vero cells were added to a 96 well plate and 24 hr later serial dilutions of 625- to 10,000-amoebae were added. Plates were incubated for 24 hr before fixing and staining with Hoechst to stain nuclei. The high content imaging assay endpoint was determined by counting the number of Vero cell nuclei remaining. (Created with BioRender.com.)

**Figure 6.**
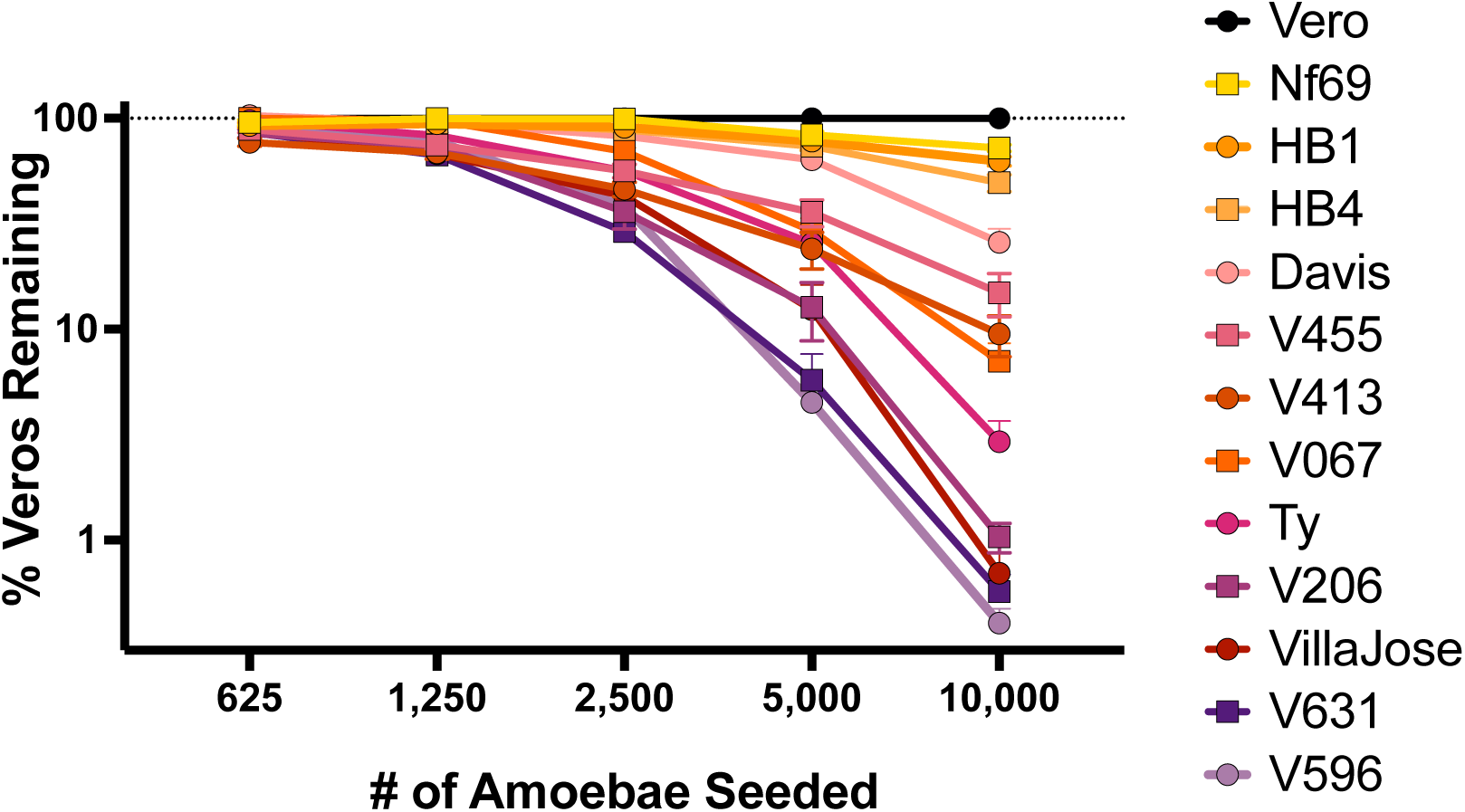
*Naegleria fowleri* isolates express varying feeding rates on Vero cells in an *in vitro* high content imaging assay. V596, V631, and Villa Jose isolates were the most virulent, whereas Nf69, HB1 and HB4 were found to be the least virulent. The assay included three biological replicates with three technical replicates for each concentration of amoebae tested.

**Figure 7.**
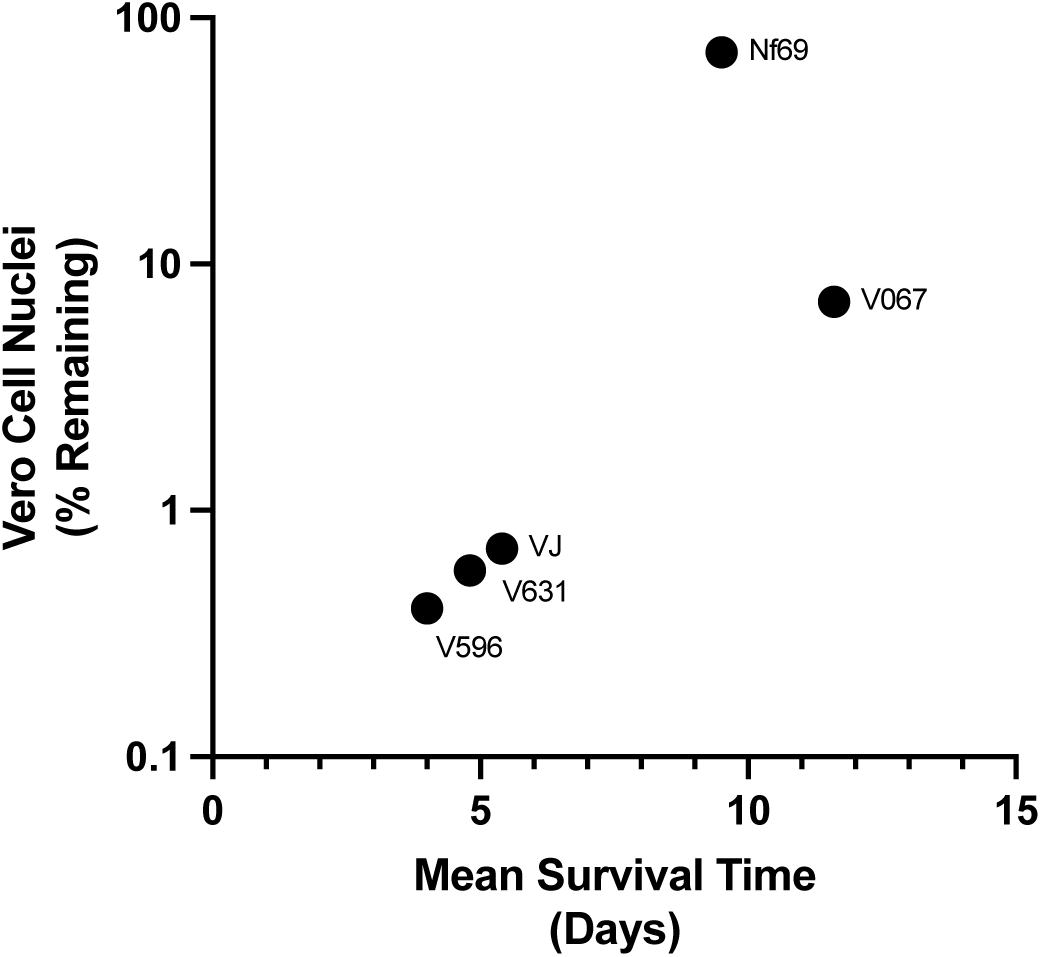
A positive correlation was observed for results of five *N. fowleri* isolates in the *in vitro* high content imaging assay and the *in vivo* mouse studies (VJ = Villa Jose).

## Discussion

Virulence of *N. fowleri* is a conundrum. There are many thermotolerant free-living amoebae in the environment, yet only *N. fowleri*, *Balamuthia mandrillaris*, *Acanthamoeba* spp. and *Sappinia pedata* are confirmed to cause disease in humans.(17, 18) There are >40 species in the *Naegleria* genus, yet only *N. fowleri* has been shown to cause a fulminant infection in humans. Furthermore, of the eukaryotic parasites known to cause disease in humans, *N. fowleri* is the most virulent given the >98% mortality rate for documented infections. Consequentially, there is significant need to better understand mechanism(s) of virulence of these amoebae and to identify virulence biomarkers that can be targeted to improve treatment outcomes of PAM patients.

In the study we aimed to determine if isolates of *N. fowleri* have innate differences in virulence. Interestingly, we observed a gradient of virulence phenotypes; there are some isolates that are highly virulent (e.g., V596), whereas there are others that are much less virulent. It is important to note that all *N. fowleri* isolates tested in this study were pathogenic (i.e., being derived from human cases and causing disease in mice) and that the major difference was the level of virulence. We also developed an *in vitro* virulence assay that positively correlates with data from the mouse model (Figure 7), plus it allows more isolates to be tested quickly and reproducibly.

A major unknown is why some people get infected with *N. fowleri* and develop PAM, whereas others with similar exposure to the same sources of amoebae contaminated water do not get infected. Many of the proposed reasons for this discrepancy involve host factors, such as immune deficiencies in the most susceptible hosts or differences in innate or adaptive immunity to amoebae.(19) In addition, it is well documented that people develop high rates of seropositivity to *N. fowleri* without ever showing signs of being infected.(5) This could be due to the route of infection *Naegleria* needs to gain access to the brain through the nasal olfactory mucosa and nerve, across the cribriform plate. Despite these data, there remains no clear answer to why PAM is a rare disease, especially in tropical climates, except for the probability it is under diagnosed or deaths are attributed to other infectious agents causing meningitis.

Several potential virulence factors have been proposed and these include a pore-forming protein, cysteine proteases, lipases, and a secreted glycosidase.(20–23) Transcriptome and comparative genomics data demonstrate the proposed virulence factors are present in both the core and accessory *Naegleria* genomes, thus suggesting that differential gene expression or *N. fowleri* specific genes of unknown function play a central role in virulence.(24) Given that all strains of *N. fowleri* appear to be pathogenic, the discovery of significant differences in virulence amongst different isolates in this study offers the possibility for comparative studies in lowly versus highly virulent strains. A recent study reported on potential virulence mechanisms,(25) yet the HB1 strain was used, and our data demonstrate HB1 is one of the least virulent strains of *N. fowleri* tested thus far (Figure 6). In another study, inocula of 50,000 amoebae (*N. fowleri* Lee strain) were used to initiate mouse infections,(26) which is 5-fold higher than required for lowly virulent strains to cause disease in this study. The large differences in amoebae inoculated, as well as the inherent virulence of the strain used in a study underscore the need for standardized methods for the study of virulence and pathogenicity mechanism(s) of *N. fowleri*.

The mouse model of PAM is most often used for virulence studies. The advantages of the mouse model include the intranasal route of infection, the acute disease progression, and the pathology induced by the amoebae, which are all similar to what is observed for PAM in humans. Despite the robust mouse model of disease, there are challenges for assessing differences in virulence between multiple amoeba isolates. First, some isolates were collected decades ago and the number of culture passages for these isolates *in vitro* are unknown. This is a problem since in this study and previously published studies, isolates grown in long term axenic culture appear less virulent than the same isolate passaged routinely in mice.(21) Secondly, virulence in mice is usually assessed as survival time following intranasal inoculation of amoebae. Although the mouse model has been widely used for drug efficacy studies,(15) there are no generally accepted standards for the isolate, the number of amoebae inoculated or the mouse strain(s) that are used. These differences result in significant variances in apparent *N. fowleri* virulence in published reports. For example, in some studies only ∼30% of infected, untreated control mice succumbed to disease and the mean survival times exceeded 14 days.(27) In contrast, in other studies 100% of the mice died by day 8.(15) The results of this study demonstrate that multiple passages of amoebae feeding on Vero cells prior to initiating mouse infections induce reproducible infections. In addition, our data demonstrate that the strain of *N. fowleri* used for *in vivo* studies requires different numbers of amoebae in the intranasal inoculation to result in 100% mortality in infected mice.

Why is discovering different levels of virulence in *N. fowleri* important? *N. fowleri* and other thermotolerant FLA are ubiquitous, yet they are not evenly distributed in water or soil.(28) In one of the few studies that quantified the number of amoebae per ml of water, estimates ranged from 35-90 amoebae/50 ml of surface water in the late summer from a pond in South Carolina, USA. (29) Similarly, in a previous study of mice swimming in water containing 105 amoebae/ml of water, approximately 70% of mice became infected.(30) Presumably a human inadvertently inhaling water during recreational activities (e.g., swimming, water skiing) might inhale approximately a ml of water; therefore, the chances for someone getting infected is more likely if the contaminated water contains highly virulent amoebae that have lower minimal infectious concentrations (e.g., V596). Thus, isolating *N. fowleri* from the environment to ascertain levels of virulence using our standardized *in vitro* virulence assay could allow for identification of virulence hot spots, allowing public health officials to issue warnings accordingly.

In this study we have developed standardized protocols to assess virulence of *N. fowleri in vitro* and *in vivo*. A key factor appears to be passaging the amoebae over Vero cells ≥5 times to best equilibrate between isolates with different passage histories. We also deduced that the rate of feeding on Vero cells *in vitro* is positively correlated with *in vivo* virulence, thus allowing new approaches for identifying and validating virulence determinants. In particular, the highly virulent V596 isolate paired with less virulent strains offer novel opportunities for comparative - omics studies to elucidate mechanism(s) of *N. fowleri* virulence.

Despite the advances in assessing virulence of *N. fowleri*, there remain many unanswered questions. We do not know if passaging multiple times over other mammalian cell lines will provide the same advantages that we found with Vero cells, though the amoebae will actively feed over other cell lines.(13) We also do not know if virulence differences will be observed in different mouse strains. We used female outbred ICR (CD-1) mice for these studies and it would be interesting to assess if V596 and lowly virulent strains (e.g., Nf69) reproduce these virulence profiles in other strains of mice, in particular inbred strains. Additionally, highly and lowly virulent *N. fowleri* strains offer an opportunity to identify potentially predisposing host factors by using different mouse strains.

In summary, herein we found that *N. fowleri* isolates demonstrated different levels of virulence in an *in vivo* model of disease. These differences in virulence were observed in both *in vivo* and *in vitro* models and were not correlated with strain genotypes. These differences may have public health implications given that lower inocula of the most virulent amoebae are able to initiate infection than previously assumed.

## Acknowledgements

This work was supported by the National Institute of Allergy and Infectious Diseases (R03AI141709 to DEK; T32AI060546-16 to ACR; T35OD010433 to ES) and the Georgia Research Alliance (to DEK).

## About the author

Antoinette Cassiopeia Russell obtained her BS in Biology at Auburn University in 2018 and earned a Ph.D. in Infectious Diseases at the University of Georgia in 2023. Her research interests are drug and biomarker discovery and host:pathogen interactions for neglected disease-causing pathogens.

