## Appendix Figures 1 and 2 for "Virulence of *Naegleria fowleri* isolates varies significantly in the mouse model of primary amoebic meningoencephalitis"

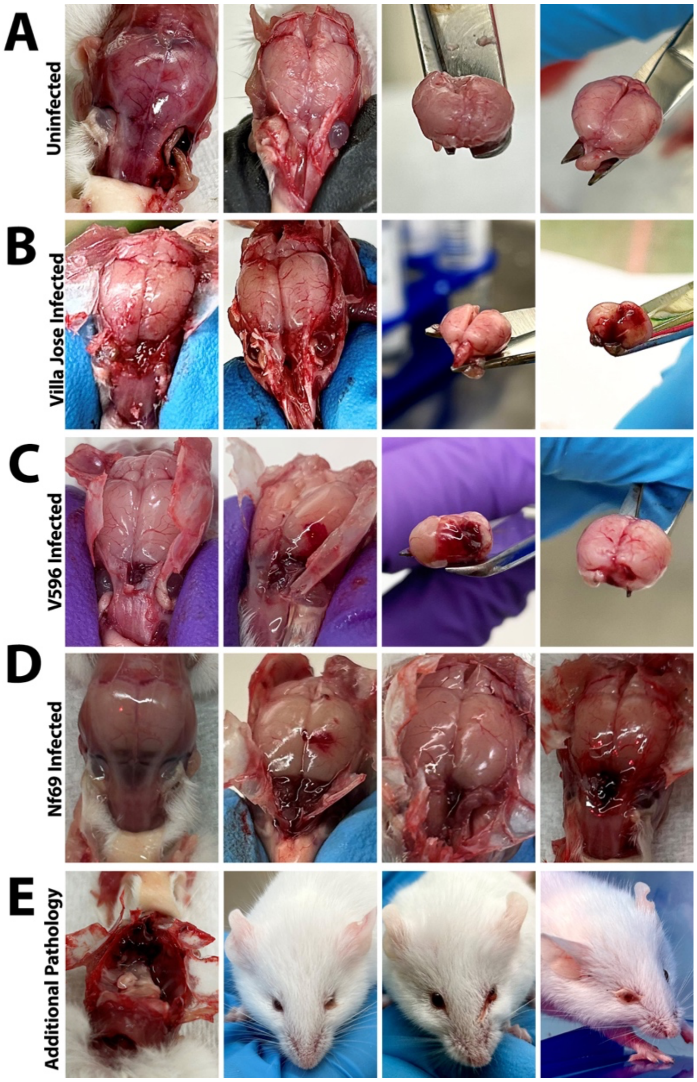


**Appendix Figure 1.** Gross pathology of brains of mice infected with different strains of *Naegleria fowleri*. Brains were dissected from mice euthanized at end stage of disease. Severe inflammation was observed in the frontal lobes of the brain in mice infected with Villa Jose (B), V596 (C), and Nf69 (D) as compared to uninfected controls (A). V596, the most virulent strain tested produced focal, punctate lesions in the olfactory bulbs, whereas the less virulent strains resulted in more diffuse pathology throughout the frontal lobes. Additional gross pathology findings (E) were liquefaction of the infected brains, eye bulging, and blood secretion from tear ducts.

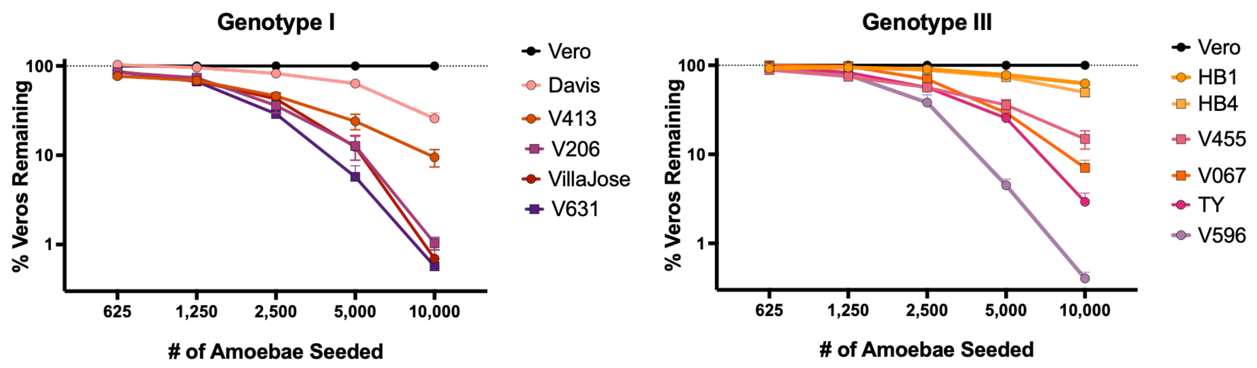


**Appendix Figure 2.** Results of the *in vitro* virulence assay demonstrated that virulence levels were not associated with genotypes I or III. The full range of virulence phenotypes, from lowly to highly virulent, were observed from both genotypes.
